# Architecture and organization of mouse posterior parietal cortex relative to extrastriate areas

**DOI:** 10.1101/361832

**Authors:** Karoline Hovde, Michele Gianatti, Menno P. Witter, Jonathan R. Whitlock

**Affiliations:** Kavli Institute for Systems Neuroscience, Norwegian University of Science and Technology, NO-7489 Trondheim, Norway

**Keywords:** cortico-cortical connectivity, tract tracing, parietal cortex, frontal cortex, prefrontal cortex, visual cortex, anatomy

## Abstract

The posterior parietal cortex (PPC) is a multifaceted region of cortex, contributing to several cognitive processes including sensorimotor integration and spatial navigation. Although recent years have seen a considerable rise in the use of rodents, particularly mice, to investigate PPC and related networks, a coherent anatomical definition of PPC in the mouse is still lacking. To address this, we delineated the mouse PPC using cyto- and chemoarchitectural markers from Nissl-, parvalbumin- and muscarinic acetylcholine receptor M2-staining. Additionally, we performed bilateral triple anterograde tracer injections in primary visual cortex (V1) and prepared flattened tangential sections from one hemisphere and coronal sections from the other, allowing us to co-register the cytoarchitectural features of PPC with V1 projections. In charting the location of extrastriate areas and the architectural features of PPC in the context of each other, we reconcile different, widely used conventions for demarcating PPC in the mouse. Furthermore, triple anterograde tracer injections in PPC showed strong projections to associative thalamic nuclei as well as higher visual areas, orbitofrontal, cingulate and secondary motor cortices. Retrograde circuit mapping with rabies virus further showed that all cortical connections were reciprocal. These combined approaches provide a coherent definition of mouse PPC that incorporates laminar architecture, extrastriate projections, thalamic, and cortico-cortical connections.

## INTRODUCTION

The posterior parietal cortex (PPC) is one of the major associational cortical areas in the brain. Across mammalian species, it receives inputs from virtually all sensory modalities, frontal motor areas and prefrontal cortex (Reep *et al.*, 1994; Krubitzer, 1995; Wise *et al.*, 1997; Stepniewska *et al.*, 2016), and it supports a variety of cognitive functions, including sensorimotor transformations, spatial processing, decision-making and movement planning. For several decades, the monkey has served as the premiere model for investigating the behavioral and neurophysiological contributions of PPC, though recent years have seen an increase in the use of rats and mice. This has been motivated in part by the fact that rodents can be trained to perform a variety of highly specific, PPC-dependent tasks in real world and virtual reality settings (Nitz, 2006; Harvey *et al.*, 2012; Raposo *et al.*, 2012; Whitlock *et al.*, 2012; Brunton *et al.*, 2013; Wilber *et al.*, 2014b; Goard *et al.*, 2016; Hwang *et al.*, 2017). The advantages of mice in particular include their genetic tractability and compatibility with large-scale recording and imaging techniques, leading to their widespread usage to study population coding and circuit function in every major sector of cortex, including PPC. Despite the popularity of the mouse for studying parietal cortex, however, there is little consensus on a coherent anatomical definition of PPC in the mouse, which is problematic because it complicates the interpretation of the wealth of new data.

As with rats, parietal cortex in the mouse is located between visual and somatosensory cortices (Paxinos & Franklin, 2012), and the existing data suggests that it has similar patterns of cortico-cortical and thalamic connectivity (Kolb & Walkey, 1987; Reep *et al.*, 1994; Harvey *et al.*, 2012; Oh *et al.*, 2014; Wilber *et al.*, 2014a; Olsen & Witter, 2016). More detailed aspects of mouse PPC anatomy, including the boundaries which distinguish it from neighboring areas, its laminar organization and chemoarchitectural profile remain ill-defined. Recent strategies for targeting PPC in mice have therefore relied either on functional mapping of extrastriate areas around PPC (Wang & Burkhalter, 2007), or on stereotactic coordinates followed by post-hoc histological comparison to one of several reference atlases (e.g. (Paxinos & Franklin, 2012; Oh *et al.*, 2014, http://connectivity.brain-map.org). These conventions may yield consistent recording locations within a study, but hamper the comparison of PPC across studies since they are based on different labeling methodologies and unrelated nomenclatures.

We sought to resolve these discrepancies by first delineating mouse PPC using cytoarchitectural and laminar criteria obtained from Nissl-, parvalbumin (PV)-, and type-2 muscarinic acetylcholine receptor (M2AChR)-immunostained coronal sections. We next performed bilateral, triple anterograde tracer injections in mouse V1 as in earlier studies (Montero, 1993; Wang & Burkhalter, 2007), and prepared flattened sections from one hemisphere and coronal sections of the other. By labeling extrastriate projections in both flattened and coronal planes, and by comparing these alongside interleaved, annotated Nissl- and M2AChR-stained coronal sections, we located the mouse PPC with respect to the major projections from V1. Based on these coordinates, we performed triple anterograde tracer injections in PPC, revealing a previously undescribed topography in parietal output to higher visual areas. Additional monosynaptic retrograde tracing with rabies virus showed that the inputs to PPC largely matched that described in rats.

## RESULTS

### Architectural features of PPC and neighboring areas

Similar to the rat, the mouse PPC lies between primary somatosensory and visual cortices, spanning approximately 600μm anterior-to-posterior, and has distinguishable medial (mPPC), lateral (lPPC) and posterior (PtP) divisions (Paxinos & Watson, 2013; Olsen & Witter 2017). To define precisely the boundaries between PPC and neighboring cortical regions, and to discern parietal sub-areas, we examined laminar architecture using Nissl staining, and chemoarchitectonic patterns using immunostaining against PV and M2AChRs.

#### Nissl staining

The PPC is bordered anteriorly by secondary motor cortex (M2), lateral to which is a narrow band of primary motor cortex (M1), and even more laterally by primary somatosensory cortex (S1), which is discernable by prominent lamination and a clearly distinguishable layer IV (Figure 1, top left). The medial PPC (mPPC) first emerges at approximately −1.55mm relative to bregma, and is apparent by its homogeneous lamination relative to the neighboring agranular retrosplenial cortex (RSA), medially, and S1, laterally. Unlike mPPC, RSA has visibly different cell densities across layers II and III, with layer V having a sparse population of large cell bodies (Figure 1, rows 1 and 2). Somatosensory cortex is discernable by a well-developed granular layer IV and clearly stratified supragranular and infragranular layers.

**Figure 1.**
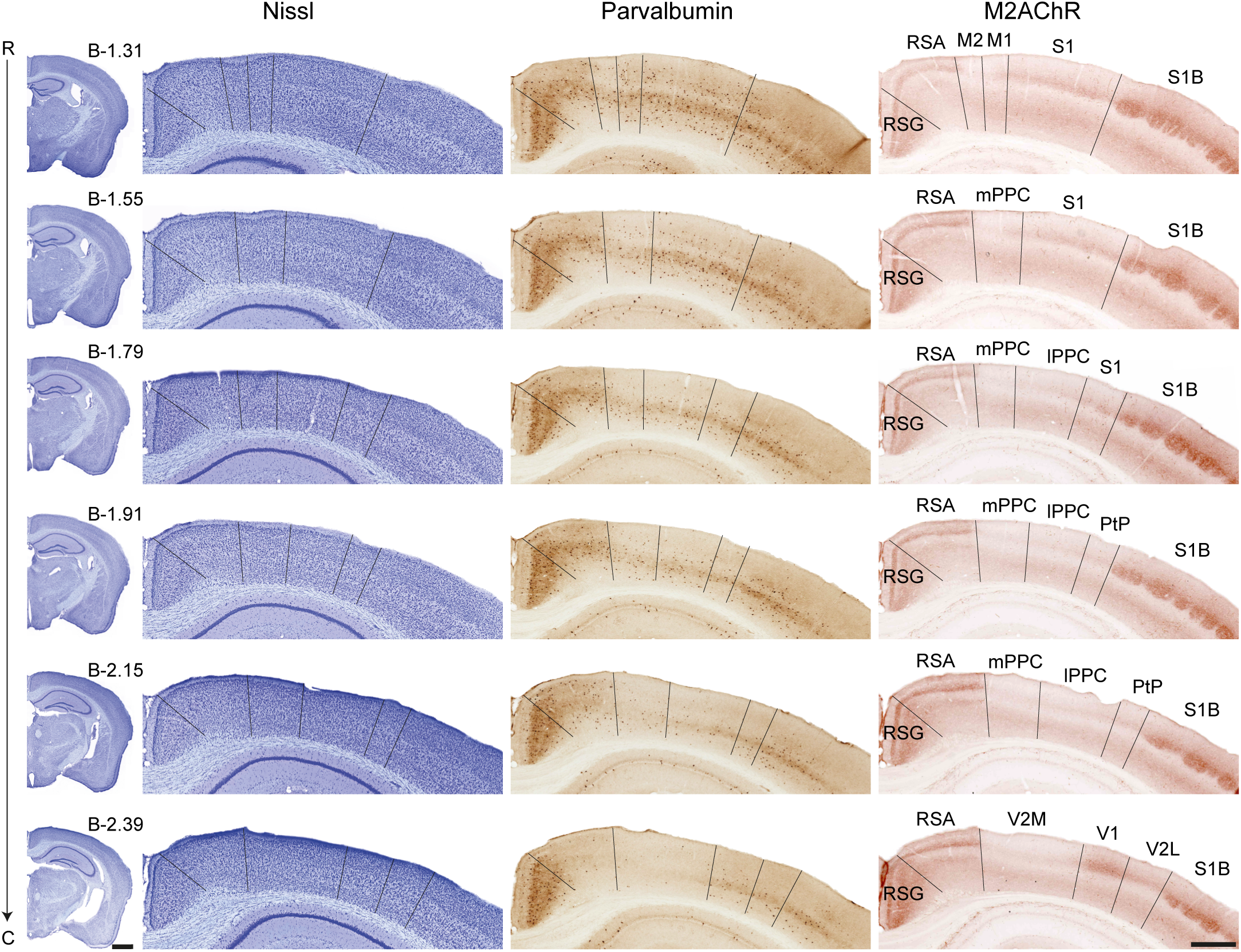
Delineation of posterior parietal cortex and surrounding areas. Coronal sections of Nissl- (left), parvalbumin- (middle) and M2AChR-stained (right) tissue from a single mouse are shown in 40μm sections in three interleaved series. Approximate bregma levels, based on Paxinos & Franklin (2012), are indicated at the far left, along with a hemispheric overview of where the sections on the right were taken from. The nomenclature is also adapted from Paxinos & Franklin (2012); see list for abbreviations. Left scale bar = 1mm, right scale bar = 500μm.

The anterior tip of lPPC appears between mPPC and S1 (Figure 1, row 3). Unlike mPPC, lPPC has some discernable lamination between layers II/III and layer V, with layer V being less densely packed than superficial layers. Layer VI of lPPC is also slightly narrower and more densely packed than mPPC. The posterior part of parietal cortex (PtP, (Paxinos & Franklin, 2012)) is the most lateral sub-area, and emerges between lPPC and S1 barrel cortex (Figure 1, row 4). Layers II/III of PtP are homogenous with small cells, whereas layer V cells are larger and more sparsely packed. Lateral to PtP is barrel cortex (Figure 1, row 5), which is distinguished by densely packed granular cells in layer IV forming the barrel fields. Caudal to mPPC and lPPC is the medial secondary visual cortex (V2M, Figure 1, row 6), which appears very similar to PPC in Nissl-staining. Considering the proximity and similarity of V2M and PPC with Nissl staining, we found that the emergence of V1, lateral to V2M, was the most useful indicator for being posterior to PPC. Primary visual cortex is characterized by a prominent, granular layer IV that is not present in V2M, and, unlike V2M, V1is clearly laminated. Similar to S1, V1 contains large pyramidal cells in layer V, and has both superficial and deep cell-sparse zones.

#### PVstaining

A salient cellular marker for the appearance of mPPC is the decrease in PV staining in both cell bodies and the neuropil (Figure 1, middle column, row 2), which contrasts with the dense staining medially in layers II-V of RSA, and the darker staining in layers IV and V of S1 (Figure 1, rows 1 and 2). The neuropil in layer V of lPPC is slightly darker than mPPC (Figure 1, row 3), though both division have distinctly less PV staining than S1 and RSA. Laterally, PtP has slightly darker staining in deep layer III and layer V than lPPC (Figure 1, row 4-5), but again it is markedly less than the neighboring S1 barrel fields, which have particularly dark staining in the neuropil of layers IV and V, and more PV+ cells in deeper layers (Figure 1, row 5). In caudal sections, V2M has even less PV staining than the neuropil and cell bodies of PPC. The near-total absence of PV staining is useful for distinguishing V2M from RSA, medially, and V1, laterally, which has a distinct band of staining in the layer V neuropil. Area V2L also has a darkly stained layer V, with weaker staining in layer superficial layers (Figure 1, row 6).

#### M2AChR stained sections

The border between RSA and mPPC is marked by a sharp decrease in M2AChR staining in the superficial parietal layers, which returns laterally in S1 (Figure 1, right column, row 2). Posterior to S1, the emergence of lPPC is also indicated by a drop in M2AChR staining (Figure 1, row 3), and again so with PtP (Figure 1, row 4). The staining in PtP is slightly darker superficially than other parietal areas, but the lateral boundary between PtP and S1B is strikingly marked by the sharp increase in superficial staining of the barrel fields. Posterior to PPC, V2M virtually lacks M2AChR staining in superficial layers, but is bracketed by strong staining in RSA and V1 (Figure 1, row 6). Here, the emergence of V1 is marked by strong M2AChR staining directly posterior to lPPC and PtP; again the appearance of V1 is the best indicator of being caudal to all PPC sub-fields in the coronal plane.

### Topography of V1 projections in tangential flattened and coronal sections

Topographic maps of V1 projections were obtained in tangential sections from flattened hemispheres containing triple-tracer injections of dextran amines at the caudal pole of V1 (Wang & Burkhalter, 2007). The tracer injections were performed bilaterally in eight mice and unilaterally in two, and a representative example of a tangential section through layer IV (Figure 2A) shows the injection sites and clusters of projections to extrastriate areas around the periphery of V1. The topography of the projections and delineations of V1 and S1 are with respect to myeloarchitectonic patterns (Figure 2, low-magnification inset) and M2AchRstaining (not shown) from the same sections.

**Figure 2.**
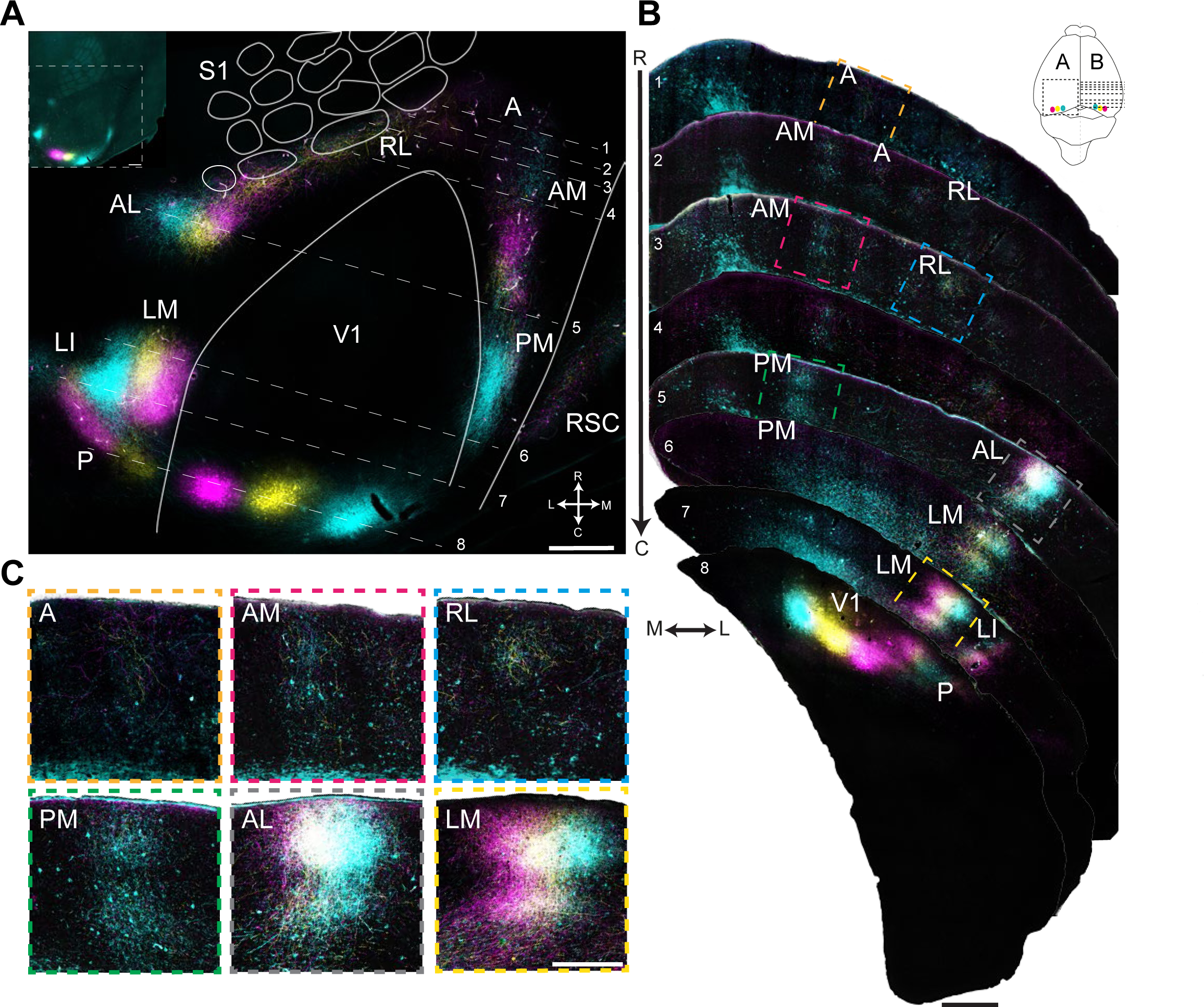
Projections from V1 viewed in flattened tangential (left) and coronal (right) sections. (**A**) Left hemisphere: Section through layer IV of flattened cortex showing triple injections of anterograde tracers in V1 and the resulting projections to extrastriate areas. Insert: dark-field image from an unprocessed section for overview. Nomenclature for the visual projection fields is based on Montero (1993). The outlines of V1 and barrels of S1 were drawn from M2AChR staining and myeloarchitecture from the dark-field image (see insert, top left). (**B**) Right hemisphere: triple injections of anterograde tracers in V1 (section 8, at bottom) as in **A**, visualized in coronal sections, as well as the resulting projections to extrastriate areas. Rostro-caudal levels of the coronal section are indicated by corresponding numbers (1 through 8) in **A**. Projections from V1 to areas LM, LI and AL were topographically organized, whereas labelling was intermingled in other sub-fields. (**C**) Magnified view of labeling from highlighted areas in coronal sections in **B**. The figure is for illustration purposes. See list for abbreviations. Scale bars in **A** and **B** = 500μm, in C = 200μm.

The projections were identified in line with previous studies (Montero, 1993; Wang & Burkhalter, 2007) based on fluorescent labeling, orientation and topographical positioning, and were named using the same nomenclature as these previous studies (see below for a nomenclatural comparison). Similar to their findings, we report a particularly strong projection from V1 to the lateromedial (LM) field, located immediately lateral to V1, and to the laterointermediate (LI) field lateral to LM (Figure 2A). Also consistent with prior observations (Wang & Burkhalter, 2007), LM showed a mirrored medial-to-lateral ordering of the labeling (Figure 2A), as did the prominently labeled anterolateral (AL) field, just posterior to the S1 barrels. The rostrolateral (RL) area contained a mixture of all tracers and ran parallel to the posterior barrel fields, while the anterior-most labeling was in the anterior (A) field, with labeling from all injections in V1 visible at higher magnifications (Figure 2C, top left). The posterior and medial to area A was the anteromedial (AM) field, with patches of labeling from each tracer stretched along the rostro-caudal axis, followed by the posteromedial (PM) field. The cyan labeling predominated at low magnification in PM, though all tracers were evident in processes at higher magnification (Figure 2C, bottom left).

Similar triple-injections were made in V1 of the right hemisphere, from which coronal sections were cut along the anterior-to-posterior extent of the extrastriate cortex (Figure 2B). We identified fields in the coronal plane based on their labeling with respect to the flat maps, and at levels corresponding to prior descriptions of the extrastriate clusters in coronal sections (D’Souza *et al.*, 2016). Consistent with the flat map, area A was labeled sparsely following injections in V1, whereas the densest projections were to areas LM and AL, both of which exhibited topographically distributed labeling. Each extrastriate area (except LI) is shown at higher magnification in Figure 2C.

### Locations of extrastriate areas in relation to PPC

A main goal of this study was to describe the location of extrastriate areas as described by Montero (1993) and Wang & Burkhalter (2007) in relation to the laminar, cyto- and chemoarchitectural features that distinguish PPC in the mouse. To do this we stacked images of Nissl-stained, annotated PPC sections atop corresponding sections from the same animal with fluorescent V1 projections, and a third series of sections stained for M2AChR (Figure 3). As shown in Figure 3B, at −1.91mm posterior to bregma the entire complement of labeled fibers for area A is contained in lPPC, with no apparent extrastriate labeling in mPPC. Proceeding caudally, area A continues to overlap with the lateral areas lPPC and PtP, while AM overlaps mainly with mPPC and to a lesser extent lPPC (Figure 3C). The bulk of labeling in AM remained in mPPC along the full extent of PPC, while area RL overlapped with PtP and the medial edge of the S1 barrel fields (Figure 3D).

**Figure 3.**
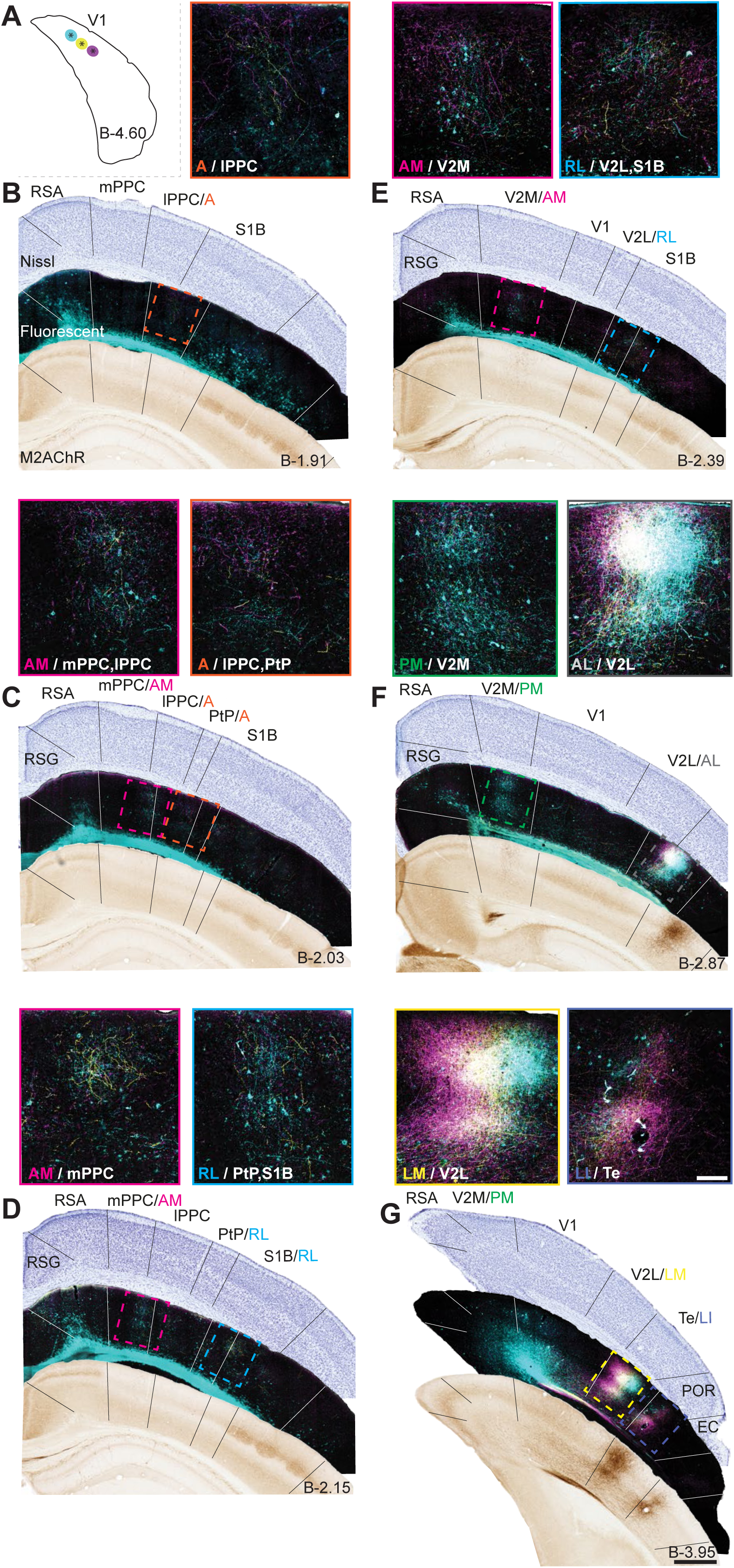
Co-registration of extrastriate areas with PPC and surrounding cortices. (**A**) Tissue from the same triple injections in V1 as in Figure 2 are shown, along with Nissl- and M2-stained sections from the same brain. Each panel consists of three immediately adjacent sections, with the series starting in PPC and proceeding caudally (AP coordinates estimated using Paxinos & Franklin, 2012). The nomenclature for extrastriate areas is based on Montero (1993) and Wang & Burkhalter (2007), and the cyto- and chemoarchitectonic labels are adapted from Paxinos & Franklin (2012). (**B**) A comparison of the three sections shows that mPPC at this level does not overlap with any extrastriate areas, whereas lPPC overlaps with area A. The enlarged inset above shows the fluorescent processes of area A / lPPC (nomenclatures juxtaposed at bottom left). (**C-G**) Similar comparisons from tissue sections spanning approximately −2 to −4mm AP. Scale bar at bottom right of **G** = 500μm; insert = 50μm.

Posterior to PPC, labelling in area AM was contained entirely in V2M (following the nomenclature of Paxinos & Franklin, 2012), and area RL continued to straddle the architectonic boundary between V2L and S1 barrels (Figure 3E). Even more caudally, at approximately −2.87mm posterior to bregma, labeling in PM overlapped completely with Paxinos & Franklin’s (2012) V2M, whereas V2L totally enveloped AL. We found that the splenium of the corpus callosum was a useful landmark for locating the transition from RL to AL, and that AL emerged at the level where the barrels disappeared in coronal sections. The farther posterior sections (Figure 3G) showed that area LM overlapped completely with caudal V2L, and area LI overlapped with the temporal association area (Te). Although we noted cross-reactivity between antibodies against M2-receptors and BDA labeling (Figures 3F-G), we confirmed that the patterns of MAChR2 labeling in other sections matched staining patterns in tissue preparations without BDA injections (as in Figure 1).

### Thalamic and cortical connectivity of mouse PPC

One of the defining features of PPC in rats and other mammals is its connection with associative thalamic nuclei, the lateral posterior (LP), lateral dorsal (LD), and posterior (Po) nucleus (McDaniel *et al.*, 1978; Donoghue & Ebner, 1981; Kolb & Walkey, 1987; Schmahmann & Pandya, 1990; Chandler *et al.*, 1992; Bucci *et al.*, 1999; Padberg & Krubitzer, 2006; Cappe *et al.*, 2007; Olsen & Witter, 2016). Existing evidence indicates that PPC in mice receives input at least from LP (Harvey *et al.*, 2012), so we used coordinates from our prior annotations (Figures 1 and 3) to target triple anterograde tracer injections in PPC (four bilaterally, and two unilaterally), and cut the right hemisphere in coronal sections to investigate the patterns of thalamic labeling. In the four bilateral cases, the left hemisphere was used to prepare tangential flattened sections.

As shown in Figures 4A and B, tracer injections contained wholly within the cytoarchitectonic boundaries of PPC produced robust anterograde labeling in LP and Po, and farther anterior sections contained strong projections confined to LD (Supplementary Figure 1). In all cases, the thalamic projections were specific to associative nuclei with no staining in the immediately adjacent dorsal lateral geniculate nucleus (DLG), which receives projections from V1, nor in the ventral posterior medial nucleus (VPM), which receives projections from S1. Thus, the cortico-thalamic projections in the mouse appear closely similar to those described in rats (Chandler *et al.*, 1992; Olsen & Witter, 2016). To visualize the position of the injection sites in PPC relative to other cortical areas, we examined flattened sections from the left hemisphere, which had similar triple injections at the same coordinates as the coronal sections. The myeloarchitecture in the flattened sections showed that our coordinates for PPC fell anterior and largely medial to V1, and tangential to the barrel fields of S1, in particular the δ barrel (Figure 4D).

**Figure 4.**
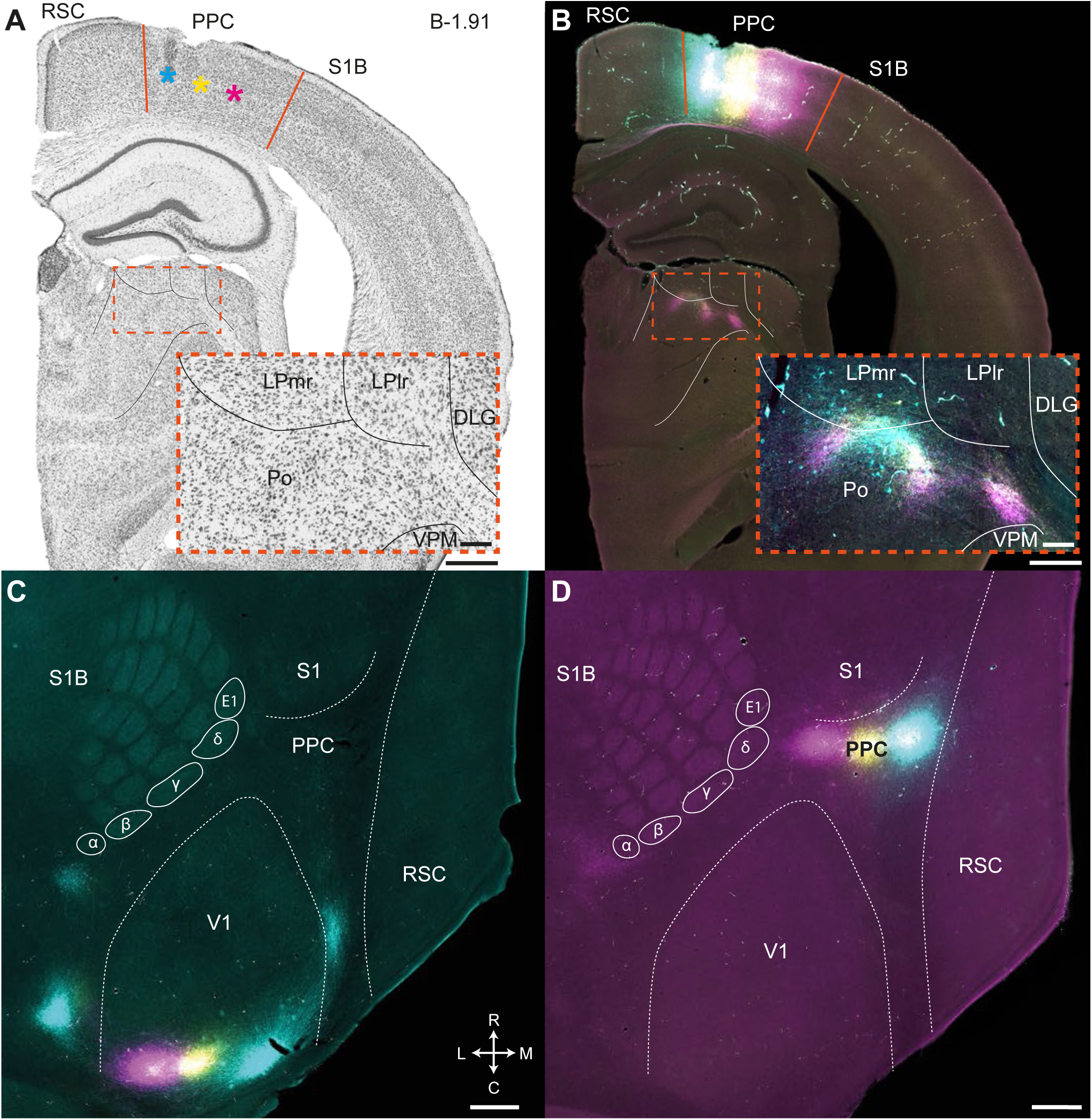
Validation of coordinates for PPC using thalamic labeling. (**A**) A Nissl-stained section from the right hemisphere showing triple injections of dextran amine tracers within the cytoarchitectonic boundaries of PPC, with the underlying thalamic nuclei delineated (inset). (**B**) Fluorescent image of the same section, showing fluorescent anterograde labeling in the associative thalamic nuclei LP and Po, with no staining in the DLG. (**C**) Overview of flattened cortex showing triple anterograde tracer injections in V1, along with S1B, S1, RSC and an estimation of where PPC should fall. (**D**) Triple anterograde injections at the same coordinates as **A**, in the left hemisphere of the same animal, viewed in a flat map with the area estimated as PPC directly lateral to the 5 barrel field. Scale bars at bottom right of **A-D** = 500μm; insert = 100μm.

To verify the projection targets of PPC, we next examined labeling resulting from triple-injections in coronal sections of the right hemisphere (Figure 5A, left), which showed prominent labeling in several cortical and sub-cortical regions. Anterior to PPC, this included projections targeting medial, ventral and ventrolateral orbitofrontal cortex (MO, VO, VLO; Figure 5B, left, MO not shown), with the most prominent labeling in superficial layers of VLO. The injections also produced strong labeling in cingulate (Cg) and secondary motor (M2) cortices, which appeared to follow a coarsely topographical distribution, with medial PPC projecting medially toward Cg, and lateral PPC projecting more laterally into M2 (Figure 5C-D, left). The projections from PPC to M2 were particularly strong posterior to bregma, though whether labeling was topographical at this level varied across animals (Supplementary Figure 2). We also noted that the projections to M2 corresponded well with outputs described from extrastriate areas A and RL (Wang *et al.*, 2012). These connections, along with robust projections to primary somatosensory cortex (S1; Figure 5E) and sub-cortical projections to the dorsal striatum and deep layers of the superior colliculus (Supplementary Figure 1), are strongly consistent with the complement of connections described in rats (Kolb & Walkey, 1987; Chandler *et al.*, 1992; Wilber *et al.*, 2014a; Olsen & Witter, 2016). To determine whether these cortical outputs of PPC were reciprocal, we performed monosynaptic circuit tracing with rabies virus in a parallel series of mice (n = 4, Figure 5A, right) (Wickersham *et al.*, 2007), which yielded unequivocal evidence that PPC received monosynaptic inputs from each cortical area with anterograde labeling (Figure 5B-E, right).

**Figure 5.**
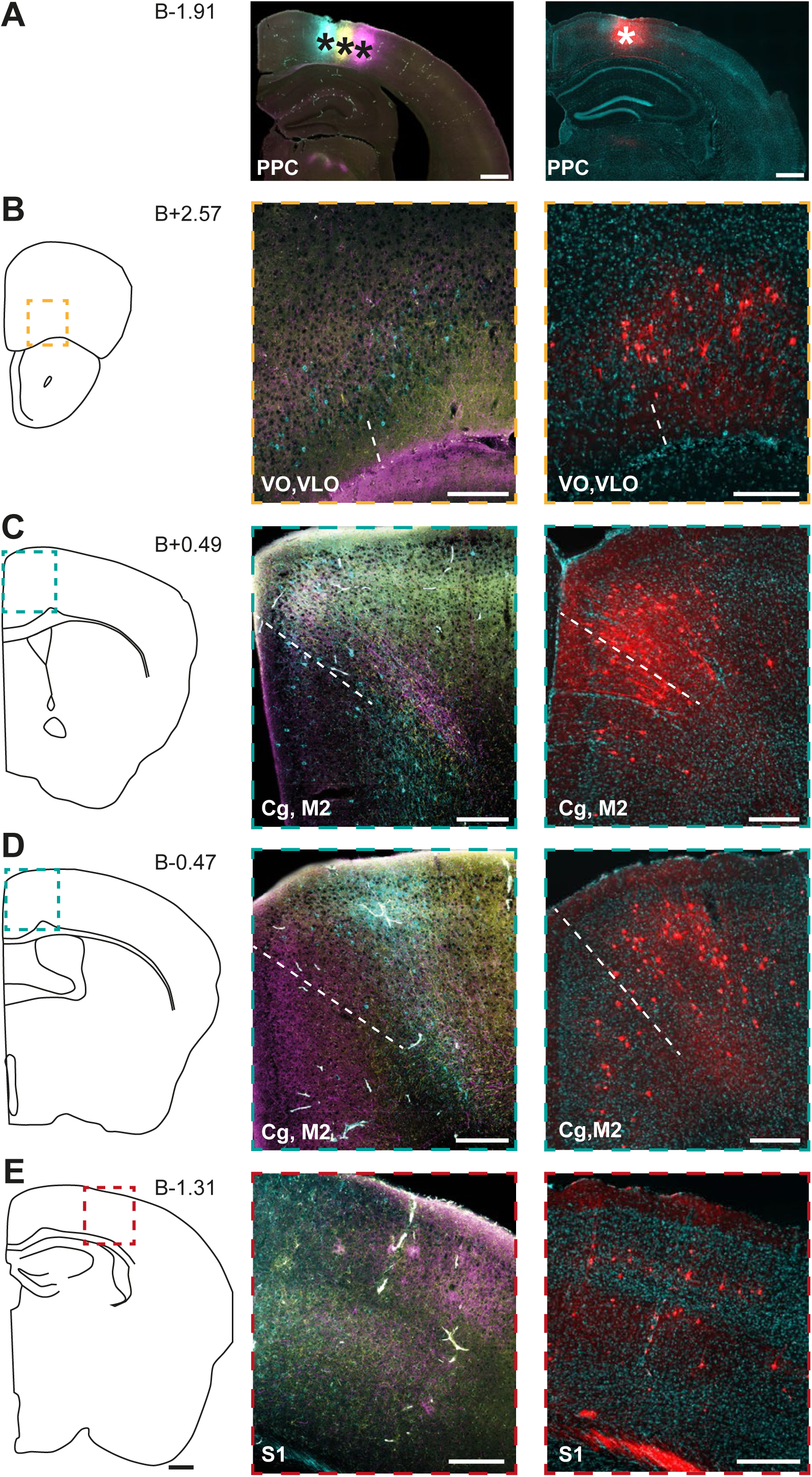
Efferent and afferent cortical connections anterior to PPC. (**A**)(left) Coronal section showing triple anterograde tracer injections in PPC of the right hemisphere; (right) injection site of TVAG and rabies viruses in the right hemisphere of a different mouse. (**B**) (left) Drawing of the right hemisphere at +2.57mm from bregma, from which the middle and right panels were taken. (middle) Fluorescent images of PPC projections to VO and VLO, (right) retrograde rabies labeling (red) against Hoechst counterstaining (blue). (**C**) (left) Drawing of the right hemisphere at +0.49mm from bregma. (middle) Strong anterograde labeling in Cg and M2, showing a rough topographical correspondence with injection sites in PPC. (right) Rabies labeling indicated dense monosynaptic projections from dorsal Cg cortex and medial M2 to PPC. (**D**) (left) Same as above, toward the caudal extent of M2, (middle) fluorescent anterograde projections from medial and lateral PPC; (right) retrogradely labeled neurons in caudal M2 and Cg that project to PPC. (**E**) (left) Drawing of the right hemisphere at −1.31mm relative to bregma. (middle) At this level, PPC has robust projections to S1, and (right) rabies labeling in S1 shows the connection is reciprocal. Scale bars in **B-E** = 500μm in left panels, 200μm in middle and right panels.

Posteriorly, PPC projections were labeled in both superficial and deep layers of primary auditory cortex (Figure 6A), and in granular and agranular retrosplenial cortex (Figure 6B, top), which is consistent with observations in rats (Kolb & Walkey, 1987; Reep *et al.*, 1994; Wilber *et al.*, 2014a; Olsen & Witter, 2016) and could correspond to RSC projections from areas A and AM in mice (Wang *et al.*, 2012; Wilber *et al.*, 2014a). The densest projections from PPC were to extrastriate areas AM/V2M and AL/V2L (Figure 6B, middle and bottom), with AM/V2M showing a medial-to-lateral topography in line with the location of tracer injections in PPC, and AL/V2L in some cases showing a mirrored ordering (Figure 6B, bottom; Supplementary Figure 2, bottom). Although labeling from PPC was present farther caudally in areas PM and LM, it was weaker than in AM/V2M and AL/V2L, and did not appear topographical (not shown). As with cortical connections anterior to PPC, monosynaptic tracing with rabies virus confirmed these connections were reciprocal, originating from both deep and superficial layers in all upstream areas (Figure 6A and B, right).

**Figure 6.**
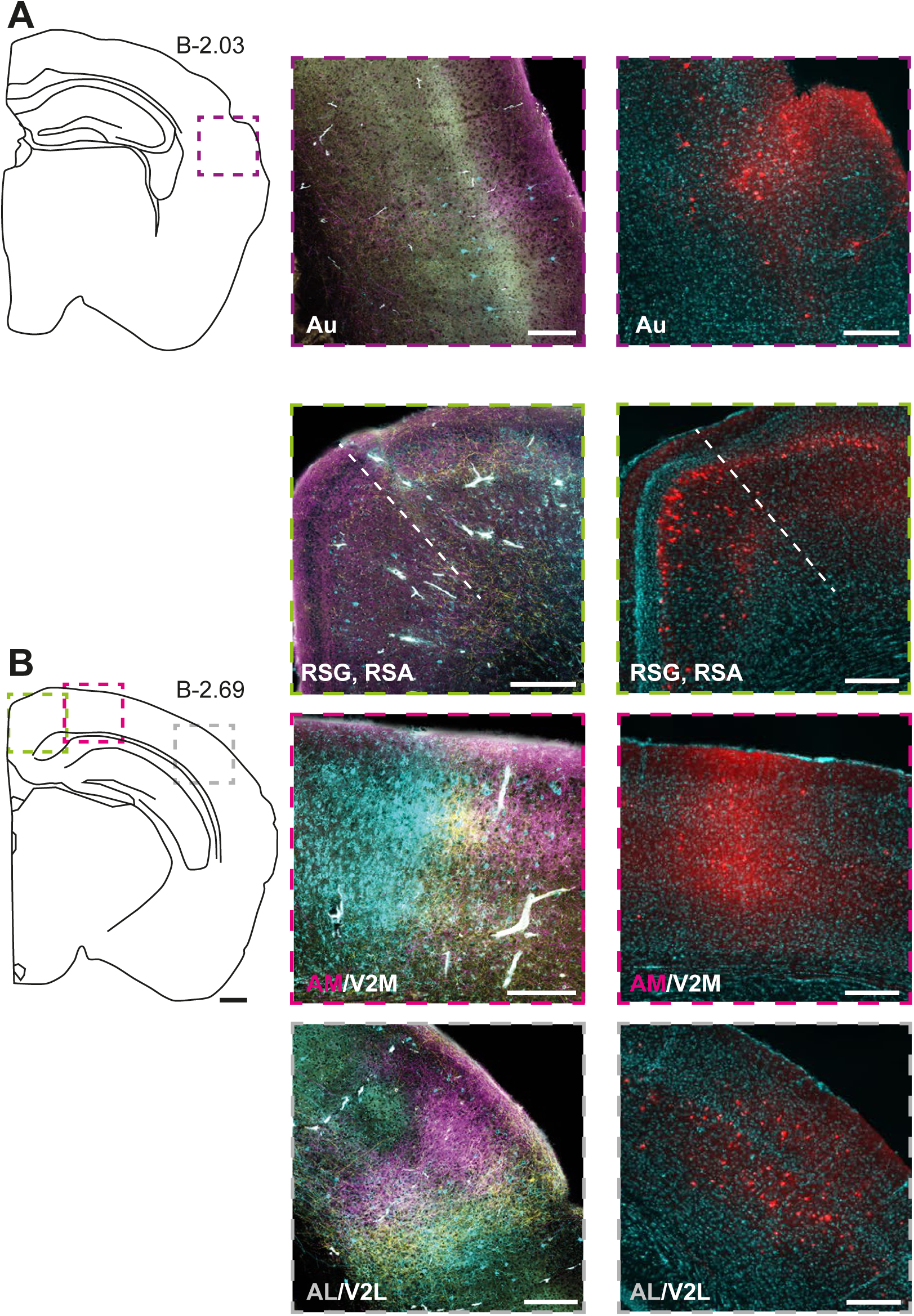
Efferent and afferent cortical connections posterior to PPC. Anterograde and retrograde labeling resulting from the same injections as in Figure 5. (**A**) (left) Drawing of the right hemisphere at −2.03mm from bregma, from which the middle and right panels were taken. (middle) PPC projects to deep and superficial layers of auditory cortex (Au); (right) Au provides monosynaptic input back to PPC, as indicated by rabies tracing. (**B**) (left) Same as above, at −2.69mm relative to bregma. (middle) PPC projections were present in superficial and deep layers of RSC (top), but were stronger and clearly topographical in AM and AL (lower panels). (right) Deep and superficial RSC, as well as areas AM and AL provide monosynaptic input to PPC. Scale bar = 500μm in left panels, 200μm in middle and right panels.

## DISCUSSION

In this study, we described laminar, cytoarchitectonic and chemoarchitectonic criteria for defining the mouse PPC and surrounding cortices, which, to our knowledge, have not been established previously. By providing a characterization of PPC and its boundaries using intrinsic architectural features, this study differs from previous, large-scale investigations of the organization of mouse cortex based on functional connectivity and projection patterns (Lim *et al.*, 2012; Oh *et al.*, 2014; Zingg *et al.*, 2014). Importantly, we reconciled widely used but disparate nomenclatures that refer to PPC (Paxinos & Franklin, 2012) *versus* the extrastriate areas around it (Montero, 1993; Wang & Burkhalter, 2007). We further confirmed our coordinates for PPC on the basis of projections to associative thalamic nuclei, which corresponded very closely to well-established thalamic projection patterns in rats (Kolb B and J Walkey 1987; Chandler HC et al. 1992; Bucci DJ et al. 1999; Olsen GM and MP Witter 2016), mice (Harvey *et al.*, 2012) as well as several other mammalian species (Donoghue & Ebner, 1981; Olson & Lawler, 1987; Schmahmann & Pandya, 1990; Padberg & Krubitzer, 2006).

Considering the growing use of mice to study PPC and the networks with which it connects, advancing a straightforward cytoarchitectonic definition of the mouse PPC in relation to nearby areas is increasingly relevant. The two major aims of this study were therefore (i) to provide a resource for identifying mouse PPC with anatomical criteria that are evident using ubiquitous staining methods, such as a Nissl stain, and (ii) to define where PPC falls in relation to extrastriate areas around V1. Early work by Montero (1993) described and named these regions in rat visual cortex, which were later characterized systematically in mice based on their anatomical inputs from V1, their retinotopic response properties (Wang & Burkhalter, 2007; Andermann *et al.*, 2011; Marshel *et al.*, 2011; Garrett *et al.*, 2014), and projections to other cortical areas (Wang *et al.*, 2012). Since the dorsal cortical surface in rodents lacks gross anatomical landmarks, the pattern of responses in extrastriate areas have provided increasingly-used functional landmarks for locating higher visual (Garrett *et al.*, 2014) and associative regions in the posterior cortex (Olcese *et al.*, 2013; Driscoll *et al.*, 2017). However, functional mapping of this kind is not feasible in the absence of a broadly expressed calcium indicator or intrinsic optical imaging, and the location of these areas relative to PPC had not been defined explicitly until now.

By characterizing the architectural boundaries of PPC and cross-referencing them with projections from V1 (Wang & Burkhalter, 2007), we established that the anterior pole of PPC does not overlap with any extrastriate areas, whereas the posterior sectors of PPC overlap partly with areas A and AM. The most lateral and posterior extents of PPC, area PtP (Paxinos & Franklin, 2012), largely overlapped with area RL. These areas are referred to as “extrastriate”, implying a primacy of visual processing (Wang & Burkhalter, 2007; Andermann *et al.*, 2011; Marshel *et al.*, 2011; Garrett *et al.*, 2014), though they overlap considerably with PPC, which has cognitive (Harvey *et al.*, 2012; Morcos & Harvey, 2016; Hwang *et al.*, 2017; Akrami *et al.*, 2018), navigational (Nitz, 2006; 2012), and movement-related (McNaughton *et al.*, 1994; Whitlock *et al.*, 2012) functions that can be expressed independently of visual input. For example, tetrode recordings in unrestrained rats targeting the caudomedial portions of PPC and extrastriate cortex (area Oc2M according to (Zilles, 1985)), which appears to coincide with area AM, showed that substantial proportions of cells were tuned to self-motion, angular head velocity and distal landmarks (Chen 1988; Chen *et al.*, 1994). Subsequent work spanning similar cortical territory also found robust coding of self-motion and landmark positions in egocentric coordinates (Wilber *et al.*, 2014b; Wilber *et al.*, 2017), again indicating roles in behavior well beyond purely visual processing, though it is possible that visual signals or optic flow contribute to such representations. The exact functions of PPC and surrounding extrastriate areas therefore merit further, systematic investigation outside of passive perceptual tasks, with substantial information likely to be gained from active or freely behaving animals.

Nevertheless, considerable portions of the mouse PPC indeed receive input from V1, and its additional connections with auditory and somatosensory areas (Zingg *et al.*, 2014) are fully consistent with a role in multisensory processing (Olcese *et al.*, 2013; Raposo *et al.*, 2014). The parietal connections with frontal cortex likely support a role in elaborating movement, whereas connections with retrosplenial cortex and the dorsal presubiculum (Zingg *et al.*, 2014; Olsen *et al.*, 2017) likely contribute to navigational functions (Whitlock *et al.*, 2008; Save & Poucet, 2009) and, possibly, transformations from first-person to third-person reference frames (Byrne *et al.*, 2007; Alexander & Nitz, 2015). The general topological relationship of PPC to these extrinsic systems has been described to various extents across species, including rats (Kolb & Walkey, 1987), cats (Olson & Lawler, 1987), ferrets (Manger *et al.*, 2002), galagos (Stepniewska *et al.*, 2016), shrews (Remple *et al.*, 2006), new world (Gharbawie *et al.*, 2011) and old world monkeys (Cavada & Goldman-Rakic, 1989), and humans (Kaas & Stepniewska, 2016). Assuming that hodological data is indicative of information flow, PPC appears to contribute to similar cognitive and behavioral functions regardless of species (Kaas, 1995; Krubitzer, 1995; Raposo *et al.*, 2012; Brunton *et al.*, 2013; Whitlock, 2014; Goldring, 2017), though the relative weight of different sensory inputs, for example, could vary according to evolutionary niches. As for mice, it remains to be mapped more fully where the inputs from associative, motor, and sensory areas are integrated synaptically within PPC, and whether graded topographies exists for different sensory modalities, as shown for visual and vibrissal afferents in area RL (Olcese *et al.*, 2013). While the present study focused on characterizing PPC relative to extrastriate projections, comparable mapping could likely be performed in the context of somatosensory, auditory or motor inputs.

## MATRIALS AND METHODS

A total number of 24 adult C57BL/6JBomTac mice (24-35 g, Taconic) were used in the current study. The mice were housed in separate cages with free access to water and food, and were kept on a reversed light-dark cycle. All surgical procedures were approved by the Norwegian Food Safety Authority as well as the local Animal Welfare Committee of the Norwegian University of Science and Technology, and followed the European Communities Council Directive and the Norwegian animal welfare act.

### Atlas brain

A 7 months old female mouse, weighing 30g, was given an overdose of pentobarbital and transcardially perfused using Ringer’s solution (0.025% KCl, 0.85% NaCl, 0.02% NaHCO3, pH 6.9) followed by a freshly prepared paraformaldehyde solution (PFA, 4% in 0.125M phosphate buffer, pH 7.4). The brain was carefully removed from the skull and post-fixed in 4% PFA overnight at 4°C. Subsequently, the brain was moved to a cryoprotective solution (2% dimethyl sulfoxide, DMSO in 0.125 M phosphate buffer, VWR) and stored again overnight at 4°C before sectioning. The brain was cut in 40μm coronal sections on a freezing microtome (Microm HM430, Thermo Scientific, Waltham, USA) in three equally spaced series. One series was used for Nissl staining, and the other two were used for immunohistochemistry against PV and M2AChR, respectively. Nissl staining and immunohistochemical procedures were the same for these and the anatomical tracing experiments, and are explained in detail in “Histology and immunochemistry” below.

### Anterograde anatomical tracing

The coordinates for initial injections were based on Paxinos and Franklin (2012), and adjusted both to the size of the animal and according to the histology of injection sites in previous animals. All surgeries were performed under isoflurane anesthesia with the animal laying on a heating pad maintaining the body temperature at 37°C. Briefly, the animal was anesthetized in a box pre-filled with isoflurane before being placed in a stereotaxic frame (Kopf Instruments). The analgesics Metacam (5 mg/kg, meloxicam, Boehringer Ingelheim Vetmedica) and Temgesic (0.1 mg/kg, buprenorphine, Indivior) were injected subcutaneously, as was the local anesthetic Marcain (1-3 mg/kg, bupivacaine, AstraZeneca) where the incision was to be made. The head of the animal was shaved, disinfected with 70% ethanol and iodine (Iodine NAF Liniment 2%, Norges Apotekerforening) and a small incision was made along the midline. The skull was cleaned with hydrogen peroxide (H_2_O_2_, 3%, Norges Apotekerforening) and 0.9% saline, the height of bregma and lambda were measured and adjusted to ensure the skull was levelled, and a craniotomy was made with a high-speed dental drill and 0.25mm burr over the coordinates for injections.

The anterograde tracers used for triple injections were (i) 10 KD biotinylated dextran amine (BDA, Dextran, Biotin, 10,000 MW, Lysine Fixable (BDA-10,000), Thermo Fisher Scientific Cat. No. D1956, RRID:AB_2307337 in 5% solution in 0.125 M phosphate buffer), (ii) Dextran, Alexa Fluor™ 488 (ThermoFisher, 10,000 MW, Anionic, Fixable, Catalog number D22910) and (iii) Dextran, Alexa Fluor™ 546 (ThermoFisher, 10,000 MW, Anionic, Fixable, Catalog number D22911). Tracers were injected iontophoretically by applying pulses of positive DC-current (6s on/off alterations, 6μA) for 10 min using glass micropipettes (20μm tip, Harvard apparatus, 30-0044). The injections were spaced 0.3 mm apart beginning 2.30 mm lateral of the midline. For all but one case, triple bilateral injections were made into left V1 and PPC. In one case, triple injections were made into V1 in the right hemisphere only. The injections into PPC were spaced 0.37mm apart, beginning 1.25mm lateral from of the midline, −1.90mm posterior to bregma. Following the injections, the craniotomy was filled with Venus Diamond Flow (Kulzer, Mitsui chemical group), the skull was cleaned with saline, and the wound was stitched and disinfected with iodine. The animals was then transferred to a heating chamber until awake and active, before being moved back to its home cage. Postoperative pain management included Metacam (5 mg/kg) 12 hours post-surgery and, if deemed necessary, 24 hours post-surgery.

### Rabies tracing

Injections were made into PPC (B-2.00, L+1.50, D-0.50) following the general surgical procedure as described above. 300nl helper virus (AAV1.CamKII0.4.Cre.SV40 + AAV5-syn-FLEX-splitTVA-EGFP-B19G, in a 1:1 ratio; Cre virus from U. Penn Vector Core; TVA virus was a generous gift from the lab of Cliff Kentros) was injected using glass capillaries (World Precision Instruments (WPI), Cat. No. 4878), a Nanoliter2010 injector (WPI) and a Nanoliter2000 pump (WPI), with the glass tip left in place 10 min after the injection. 12 days later, 230nl of rabies virus (EnvA-pseudotyped SAD-DeltaG-mCherry; gift from Kentros lab) was injected in the same location, and the animal was kept alive for 11 days before perfusion.

### Tissue collection and preparation

One week after tracer injections, the animal was given an overdose of pentobarbital and perfused transcardially with Ringer’s solution (0.025% KCl, 0.85% NaCl, 0.02% NaHCO3, pH 6.9) followed by freshly prepared paraformaldehyde solution (PFA, Sigma-Aldrich AS, 1% in 0.125M phosphate buffer, pH 7.4). The brain was carefully removed from the skull and kept in a container with PFA (1%). Within one hour of the perfusion, the brain was cut in two along the midline to prepare coronal sections of the right hemisphere and tangential sections through flattened tissue (flat maps) from the left hemisphere.

### Coronal sections

The right hemisphere was transferred to a screw-top vial containing PFA (4%, 0.125M phosphate buffer, pH 7.4), post-fixed overnight at 4°C and transferred to cryoprotective solution (2% DMSO in 0.125 M phosphate buffer) the next day, and again stored overnight at 4°C. The hemisphere was then cut in 40μm coronal sections on a freezing microtome (see above) in three equally spaced series. The first series was mounted directly onto Superfrost Plus microscope slides (Gergard Menzel GmbH, Braunschweig, Germany), dried overnight on a heating pad, and used for Nissl staining. Series two and three were each stored in cryoprotective solution at −20°C, and later used for visualizing anterograde tracers or rabies, and the other for immunohistochemistry against M2AChR.

### Tangential Flattened sections

The left hemisphere was flattened to make “flat maps” of cortex, which first required that cortex was dissected from the rest of the subcortical structures. This was done by first resting the left hemisphere on the midline with the cortex upwards, and gently pressing the cortex flat. The brain was next flipped over to expose the midline, and a cut was made in fornix dorsal to the anterior commissure. Two brushes were used to push and separate cerebellum, cortex and the underlying subcortical areas from each other. The brainstem was held down with a brush, while dorsal cortex and hippocampus were pushed away with dissection scissors, and cuts were made at the same time along the white matter. The scissors were held parallel to the cutting plane and special care was taken to not damage ventral hippocampus. The brainstem and cerebellum were cut out and removed. One relief cut was made in cingulate cortex and one ventral to postrhinal cortex to facilitate the unfolding of cortex. The cortex was then placed on a microscope glass covered with parafilm (Laboratory film, Pechiney, plastic Packaging, Chicago), and hippocampus and dorsal cortex were gently unfolded using two brushes. Another covered microscope glass was placed on top of the tissue and the two glasses were taped together. The preparation was placed in a container with PFA (4%) overnight at 4°C with a glass weight (52 g) on top to provide extra pressure. The following day, the flattened tissue was removed from the microscope glasses and kept in a screw-top container with 2% DMSO in 0.125 M phosphate buffer overnight at 4°C. The following day the brain was mounted onto a freezing microtome stage using a sucrose solution (20%) with dorsal cortex facing down, and 50μm thick sections of flattened cortex were cut and collected in one tube containing 2% DMSO in 0.125 M phosphate buffer. The sections were first used for studying projections from V1 to extrastriate areas in flat maps, and were stained subsequently with DAB against M2AChR for delineation purposes (as described in the following section).

## Histology and immunochemistry

### Nissl staining

Series one from the right hemisphere was stained with cresyl violet (Sigma-Aldrich, St. Louis, MO). Briefly, the sections were dehydrated in increasing percentages of ethanol (50%, 70%, 80%, 90%, 3 × 100%, 10 dips each), cleared in xylene for 2 min, and rehydrated in decreasing concentration of ethanol. The sections were rinsed briefly in running water before being stained with cresyl violet (0.1%) on a shaker for 3 min. The sections were rinsed subsequently in running water and differentiated in an ethanol-acetic acid solution (0.5% acetic acid in 70% ethanol) until optimal staining was achieved. The sections were again dehydrated in increasing percentages of ethanol (as described above), cleared in xylene, and coverslipped with entellan (Merck KGaA, Darmstadt, Germany).

### Immunohistochemistry against BDA

Series two of the right hemisphere and all sections of the left hemisphere were used for triple-anterograde tracing experiments. Two of the tracers were conjugated with Alexa fluorophores Dextran, Alexa Fluor™ 488 and Dextran, Alexa Fluor™ 546, whereas BDA was visualized using fluorophore-tagged streptavidin (Thermo Fisher Scientific). This was done by first washing tissue sections 3 * 5 min in 0.125M phosphate buffer (pH 7.4), followed by 3 × 5 min in TBS-T× (0.5% Triton-X-100, 0.606% Tris(hydroxymethyl)aminomethane, 0.896% NaCl, pH 8.0). The sections were then incubated with primary antibody Streptavidin, Alexa Fluor 633 conjugate (1:400, Thermo Fisher Scientific Cat. No. S-21375, RRID:AB_2313500) in TBS-Tx for 90 min at room temperature, followed by 3 × 5 min rinsing in Tris buffer 0.606% (Tris(hydroxymethyl)aminomethane, pH 7.6). The tissue sections were then mounted on Menzel-glass slides (Thermo Scientific) using a Tris-gelatin solution (0.2% gelatin in Tris-buffer, pH 7.6), air dried overnight and coverslipped with an entellan-toluene solution the following day.

### DAB staining against M2AChR and PV

Tissue sections were stained with 3.3’-Diaminobenzidine tetrahydrochloride (DAB, Sigma-Aldrich, St. Louis, USA) to visualize M2AChR density in series three of the right hemisphere and flat map sections for the anterograde tracer experiments, as well as for series three of the atlas brain (Figure 1). DAB staining was also used to visualize PV in series two of the atlas brain. The staining procedure for coronal sections was the same across experiments except for flat maps, for which staining was done on the slide, requiring a longer incubation time.

In brief, for immunostaining against M2AChR and PV, sections were first rinsed 2 × 5 min in phosphate buffer (0.125M) followed by 2 × 5 min rinses in TBS-Tx. The sections were incubated with primary antibody (Rat anti-muscarinic acetylcholine receptor M2 monoclonal antibody, unconjugated, clone m2-2-b3, 1:750, Millipore Cat. No. MAB367, RRID:AB_94952; Mouse anti-parvalbumin monoclonal antibody, unconjugated, clone PARV-19, 1:1000, Sigma-Aldrich Cat. No. P3088, RRID:AB_477329) overnight at room temperature. They were then washed 2 × 5 min in TBS-Tx and incubated with mouse absorbed, rabbit-anti-rat secondary antibody (Anti-rat IgG (H+L), 1:300, Vector Laboratories Cat. No. BA-4001, RRID:AB_10015300; Goat anti-mouse IgG, biotin conjugated, 1:200, Sigma-Aldrich Cat. No. B7151, RRID:AB_258604) for 90 min at room temperature. The sections were then washed 2 × 5 min in TBS-Tx, 2 × 5 min in PB, 2 × 5 min in H_2_O_2_-metanol solution (0.08%, Sigma-Aldrich), 2 × 5 min TBS-Tx and incubated with a Vector ABC kit (Vector laboratories, Inc., Burlingame, USA) for 90 min at room temperature, per the manufacturer’s instructions. Subsequently, the sections were washed 2 × 5 min in TBS-Tx, then 2 × 5 min in Tris-buffer before being incubated with DAB (10 mg in 15mL Tris-buffer, Sigma-Aldrich) at room temperature. Just before the incubation, H_2_O_2_ (2μL, 30%, Sigma-Aldrich) was added to the DAB solution and it was filtered. The sections were incubated in DAB until they reached the desired color, rinsed in Tris-buffer solution and mounted on Menzel glass slides using a 0.2% gelatin solution. After drying overnight, the slides were coverslipped with an entellan-xylene solution.

### Immunohistochemistry against rabies

For brains used for rabies tracing, series two was stained against green fluorescent protein (GFP) and red fluorescent protein (RFP) to visualize the helper virus (AAV1 .CamKII0.4.Cre.SV40 + AAV5-syn-FLEX-splitTVA-EGFP-B19G) and the rabies virus (EnvA-pseudotyped SAD-DeltaG-mCherry), respectively. In brief, the tissue was rinsed 3 × 5 min in phosphate buffered saline (PBS, 0.1M, pH 7.4, Sigma-Aldrich) on a shaker at room temperature and rinsed 2 × 10 min in a 0.3% Triton solution (PBS 0.1M and 0.3% Triton). Further, it was incubated with primary antibodies (Rabbit RFP Antibody Pre-adsorbed, 1:1000, Rockland Cat. No. 600-401-379, RRID:AB_2209751; Chicken anti-GFP, 1:500, Abcam Cat. No. ab13970, RRID:AB_300798) in a PBS 0.1M + 0.3% Triton + 3% BSA solution on a shaker at 4°C overnight. The tissue was rinsed 2 × 5 min in 0.3% Triton solution and incubated with secondary antibodies (F(ab)2-goat antirabbit IgG (H+L) cross-adsorbed, Alexa Fluor 546, 1:1000, Thermo Fisher Scientific Cat. No. A-11071, RRID:AB_2534115; Goat anti-chicken IgY H&L, Alexa Fluor^®^ 488, 1:1000, Abcam Cat# ab150169, RRID:AB_2636803) in a PBS 0.1M + 0.3% Triton + 3% BSA solution on a shaker at room temperature for one hour. Finally, the tissue was rinsed 2 × 10 min in PBS (0.1M) and mounted on gelatin-coated polysine slides (Thermo Scientific) using PBS (0.1M). After drying for one hour, a Hoechst solution (1:5000 in PBS 0.1M, bisBenzimid H 33258, catalog No. B1155, Sigma-Aldrich) was applied on the sections for 5 min in the dark, the slides were carefully rinsed with PBS and coverslipped with ProLong^®^ Gold antifade reagent (REF P36934, Molecular probes, Life technologies™).

## Imaging and analysis

All brain sections were digitized using a Zeiss Axio Scan.Z1 scanner. Selected fluorescent coronal sections and fluorescent flat maps were scanned with a Zeiss confocal microscope (LSM800). The scans were edited in Adobe Photoshop CC 2017 and figures were made in Adobe Illustrator CC 2017. Nissl and DAB stained sections were optimized for brightness and contrast. Fluorescent flat maps and coronal sections in Figures 2, 3, 4, 5 and 6 were used for illustration purposes, and were optimized in Adobe Photoshop for brightness, contrast, levels and by applying masks. This was done to reduce background and enhance visualization of labeling.

## ACKNOWLEDGEMENTS

This work was supported by an ERC starting grant (agreement N° 335328) to J.R.W., a Research Council of Norway grant to J.R.W. (agreement N° 239963), the Centre of Excellence scheme of the Research Council of Norway (Centre for Neural Computation, grant N° 223262) and the National Infrastructure scheme of the Research Council of Norway - NORBRAIN, grant N° 197467. We thank G. M. Olsen for technical assistance, aiding with annotation and helpful discussions on comparative labeling in rats and mice; A. Burkhalter for generously sharing the flat map protocol; R.R. Nair and C. Kentros for generously sharing TVA and rabies viruses; H. Kleven and H. Waade for technical and IT assistance; members of the Whitlock lab for helpful discussions.

## AUTHOR CONTRIBUTIONS

JRW and MPW conceived of and designed the study. KH performed all surgeries and histology. KH and MG analyzed the data. JRW and KH drafted the manuscript with input from MPW and MG.

## DATA ACCESSIBILITY

Scans and confocal images of tissue used in this study can be made available upon request.

## ABBREVIATIONS

### Cortical areas

Au: Auditory cortex
Cg: Cingulate cortex
EC: Entorhinal cortex
lPPC: Lateral posterior parietal cortex
M1: Primary motor cortex
M2: Secondary motor cortex
MO: Medial orbitofrontal cortex
mPPC: Medial posterior parietal cortex
POR: Postrhinal cortex
PPC: Posterior parietal cortex
PtP: Posterior part of parietal cortex
RSA: Agranular retrosplenial cortex
RSC: Retrosplenial cortex
RSG: Granular retrosplenial cortex
S1: Primary somatosensory cortex
S1B: Barrel fields of primary somatosensory cortex
Te: Temporal association cortex
V1: Primary visual cortex
V2L: Lateral secondary visual cortex
V2M: Medial secondary visual cortex
VLO: Ventrolateral orbitofrontal cortex
VO: Ventral orbitofrontal cortex

**Extrastriate areas**
A: Anterior area
AL: Anterolateral area
AM: Anteromedial area
LI: Laterointermediate area
LM: Lateromedial area
MM: Mediomedial area
P: Posterior area
PM: Posteriomedial area
RL: Rostrolateral area

**Subcortical areas**
DLG: Dorsal lateral geniculate nucleus
LD: Laterodorsal nucleus
LP: Lateral posterior nucleus
LPlr: Lateral posterior nucleus, rostrolateral part
LPmr: Lateral posterior nucleus, rostromedial part
Po: Posterior thalamic nuclear group
SC: Superior colliculus
VPM: Ventral posterior nucleus, medial part

**Visualized protein**
M2AChR: Muscarinic acetylcholine receptor type 2
PV: Parvalbumin

## Tracers, viruses and antibodies used in the study

- DA488: Dextran, Alexa Fluor™ 488; 10,000 MW, Anionic, Fixable, Thermo Fisher, Cat. No. D22910
- DA546: Dextran, Alexa Fluor™ 546; 10,000 MW, Anionic, Fixable, Thermo Fisher, Cat. No. D22911
- BDA: Dextran, Biotin, 10,000 MW, Lysine Fixable (BDA-10,000), Thermo Fisher Scientific Cat. No. D1956, RRID:AB_2307337
- AAV1.CamKII0.4.Cre.SV40, U. Penn Vector Core, Perelman School of Medicine, University of Pennsylvania
- AAV5-syn-FLEX-splitTVA-EGFP-B19G (Cliff Kentros lab)
- EnvA-pseudotyped SAD-DeltaG-mCherry (Cliff Kentros lab)
- Streptavidin, Alexa Fluor 633 conjugate, 1:400, Thermo Fisher Scientific Cat. No. S-21375, RRID:AB_2313500
- Rat anti-muscarinic acetylcholine receptor m2 monoclonal antibody, unconjugated, clone m2-2-b3, 1:750, Millipore Cat. No. MAB367, RRID:AB_94952
- Mouse anti-parvalbumin monoclonal antibody, unconjugated, clone PARV-19, 1:1000, Sigma-Aldrich Cat. No. P3088, RRID:AB_477329
- Anti-rat IgG (H+L), mouse adsorbed, made in rabbit antibody, 1:300, Vector Laboratories Cat. No. BA-4001, RRID:AB_10015300
- Goat anti-mouse IgG, biotin conjugated, 1:200, Sigma-Aldrich Cat# B7151, RRID:AB_258604
- Rabbit RFP Antibody Pre-adsorbed, 1:1000, Rockland Cat. No. 600-401-379, RRID:AB_2209751
- Chicken anti-GFP, 1:500, Abcam Cat. No. ab13970, RRID:AB_300798
- F(ab)2-goat anti-rabbit IgG (H+L) cross-adsorbed, Alexa Fluor 546, 1:1000, Thermo Fisher Scientific Cat. No. A-11071, RRID:AB_2534115
- Goat anti-chicken IgY H&L, Alexa Fluor^®^ 488, 1:1000, Abcam Cat. No. ab150169, RRID:AB_2636803

**Supplementary Figure 1.**
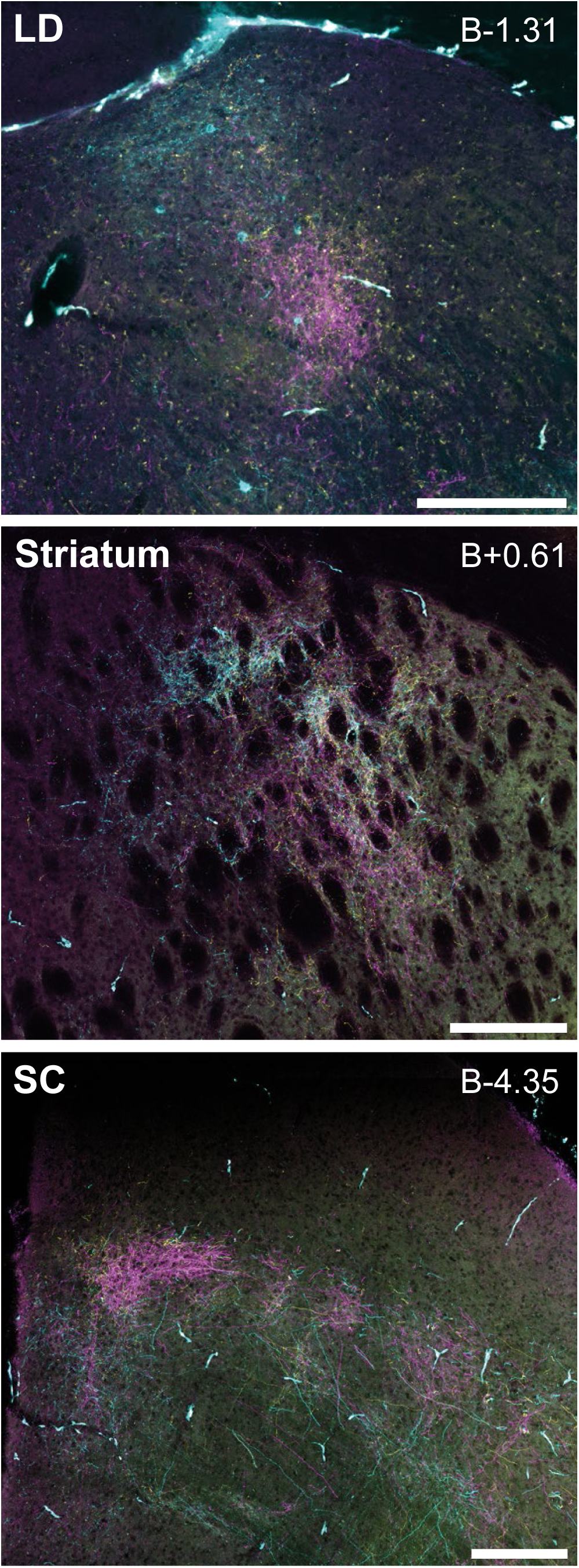
Subcortical targets of PPC projections. (top) Anterograde labeling in the lateral dorsal (LD) nucleus of the thalamus was readily apparent following triple anterograde tracer injections in PPC. This section is from the same brain as in Figures 4A and B, anterior to thalamic labeling in LP and Po. (middle) Projections to the dorsal striatum and (bottom) intermediate layers of the superior colliculus. Scale bars = 200μm.

**Supplementary Figure 2.**
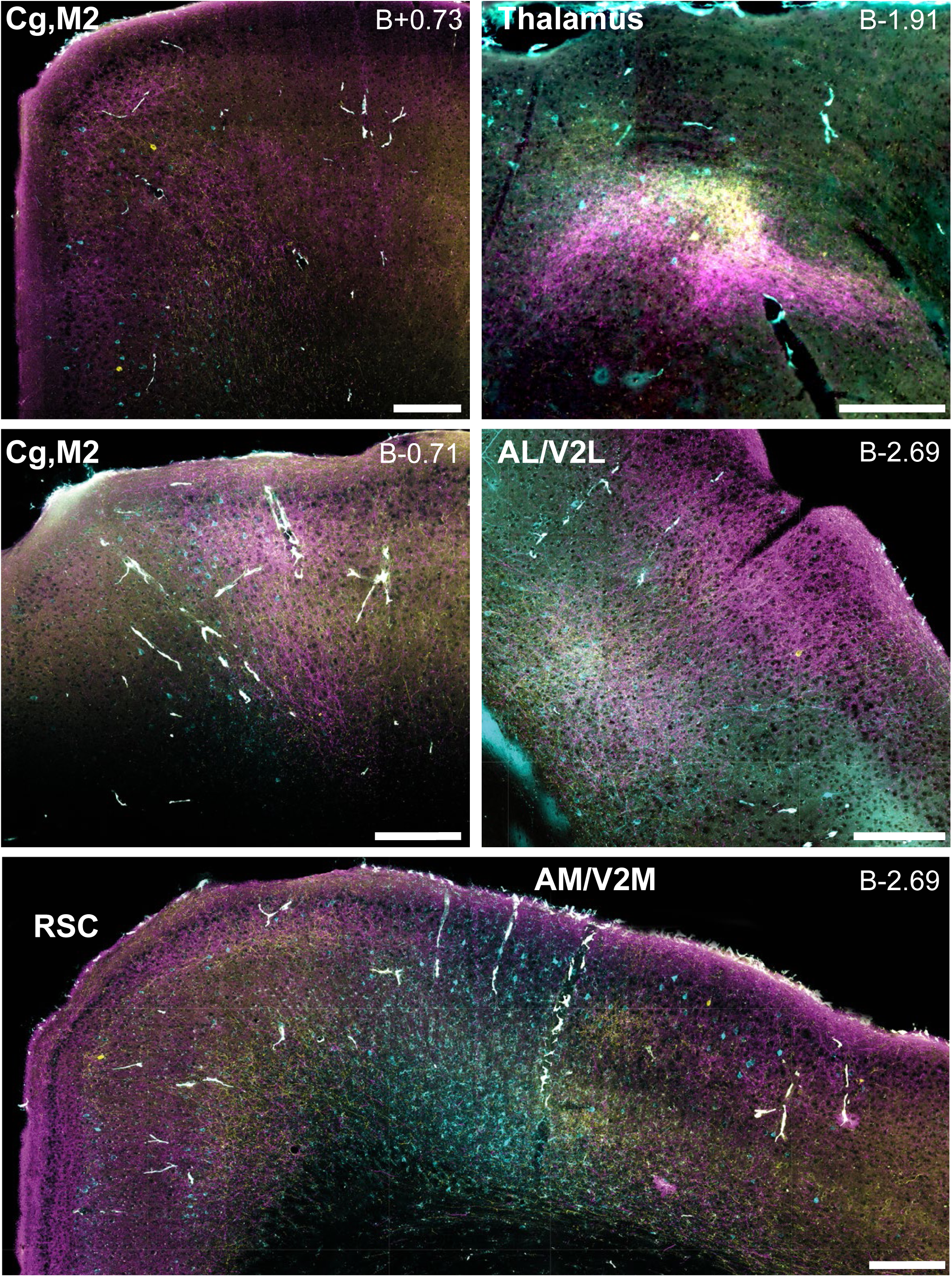
Projections of PPC were consistent across mice. Anterograde labeling following triple tracer injections in another mouse closely matched the patterns shown in Figures 4, 5, 6. This includes labeling in rostral and caudal Cg/M2 (top left panels), the LP and Po nuclei of the thalamus (top right), area AL (middle right), as well as all layers of RSC and AM (bottom). Scale bars = 200μm.

## REFERENCES

Akrami, A., Kopec, C.D., Diamond, M.E. & Brody, C.D. (2018) Posterior parietal cortex represents sensory history and mediates its effects on behaviour. Nature, 554, 368–372.

Alexander, A.S. & Nitz, D.A. (2015) Retrosplenial cortex maps the conjunction of internal and external spaces. Nature neuroscience, 18, 1143–1151.

Andermann, M.L., Kerlin, A.M., Roumis, D.K., Glickfeld, L.L. & Reid, R.C. (2011) Functional specialization of mouse higher visual cortical areas. Neuron, 72, 1025–1039.

Brunton, B.W., Botvinick, M.M. & Brody, C.D. (2013) Rats and humans can optimally accumulate evidence for decision-making. Science (New York, N.Y, 340, 95–98.

Bucci, D.J., Conley, M. & Gallagher, M. (1999) Thalamic and basal forebrain cholinergic connections of the rat posterior parietal cortex. Neuroreport, 10, 941–945.

Byrne, P., Becker, S. & Burgess, N. (2007) Remembering the past and imagining the future: a neural model of spatial memory and imagery. Psychol Rev, 114, 340–375.

Cappe, C., Morel, A. & Rouiller, E.M. (2007) Thalamocortical and the dual pattern of corticothalamic projections of the posterior parietal cortex in macaque monkeys. Neuroscience, 146, 1371–1387.

Cavada, C. & Goldman-Rakic, P.S. (1989) Posterior parietal cortex in rhesus monkey: I. Parcellation of areas based on distinctive limbic and sensory corticocortical connections. The Journal of comparative neurology, 287, 393–421.

Chandler, H.C., King, V., Corwin, J.V. & Reep, R.L. (1992) Thalamocortical connections of rat posterior parietal cortex. Neuroscience letters, 31, 237–242.

Chen, L.L., Lin, L.H., Barnes, C.A. & McNaughton, B.L. (1994) Head-direction cells in the rat posterior cortex. ii. Contributions of visual and ideothetic information to the directional firing. Experimental brain research. Experimentelle Hirnforschung, 101, 24–34.

Chen, L.L.M., B. L. (year) Spatially selective discharge of vision and movement modulated posterior parietal neurons in the rat. Vol. 14, Soc. Neurosci. Abstracts. City. p. 818.

D’Souza, R.D., Meier, A.M., Bista, P., Wang, Q. & Burkhalter, A. (2016) Recruitment of inhibition and excitation across mouse visual cortex depends on the hierarchy of interconnecting areas. Elife, 5.

Donoghue, J.P. & Ebner, F.F. (1981) The organization of thalamic projections to the parietal cortex of the Virginia opossum. The Journal of comparative neurology, 198, 365–388.

Driscoll, L.N., Pettit, N.L., Minderer, M., Chettih, S.N. & Harvey, C.D. (2017) Dynamic Reorganization of Neuronal Activity Patterns in Parietal Cortex. Cell, 170, 986–999 e916.

Garrett, M.E., Nauhaus, I., Marshel, J.H. & Callaway, E.M. (2014) Topography and areal organization of mouse visual cortex. J Neurosci, 34, 12587–12600.

Gharbawie, O.A., Stepniewska, I. & Kaas, J.H. (2011) Cortical connections of functional zones in posterior parietal cortex and frontal cortex motor regions in new world monkeys. Cereb Cortex, 21, 1981–2002.

Goard, M.J., Pho, G.N., Woodson, J. & Sur, M. (2016) Distinct roles of visual, parietal, and frontal motor cortices in memory-guided sensorimotor decisions. Elife, 5.

Goldring, A., Krubitzer, L. (2017) Evolution of parietal cortex in mammals: From manipulation to tool use. In Krubitzer, L., Kaas, J.H. (eds) The Evolution of Nervous Systems. Elsevier, London, pp. 259–286.

Harvey, C.D., Coen, P. & Tank, D.W. (2012) Choice-specific sequences in parietal cortex during a virtual-navigation decision task. Nature, 484, 62–68.

Hwang, E.J., Dahlen, J.E., Mukundan, M. & Komiyama, T. (2017) History-based action selection bias in posterior parietal cortex. Nat Commun, 8, 1242.

Kaas, J.H. (1995) The evolution of isocortex. Brain Behav Evol, 46, 187–196.

Kaas, J.H. & Stepniewska, I. (2016) Evolution of posterior parietal cortex and parietal-frontal networks for specific actions in primates. The Journal of comparative neurology, 524, 595–608.

Kolb, B. & Walkey, J. (1987) Behavioural and anatomical studies of the posterior parietal cortex in the rat. Behavioural brain research, 23, 127–145.

Krubitzer, L. (1995) The organization of neocortex in mammals: are species differences really so different? Trends Neurosci, 18, 408–417.

Lim, D.H., Mohajerani, M.H., Ledue, J., Boyd, J., Chen, S. & Murphy, T.H. (2012) In vivo Large-Scale Cortical Mapping Using Channelrhodopsin-2 Stimulation in Transgenic Mice Reveals Asymmetric and Reciprocal Relationships between Cortical Areas. Front Neural Circuits, 6, 11.

Manger, P.R., Masiello, I. & Innocenti, G.M. (2002) Areal organization of the posterior parietal cortex of the ferret (Mustela putorius). Cereb Cortex, 12, 1280–1297.

Marshel, J.H., Garrett, M.E., Nauhaus, I. & Callaway, E.M. (2011) Functional specialization of seven mouse visual cortical areas. Neuron, 72, 1040–1054.

McDaniel, W.F., McDaniel, S.E. & Thomas, R.K. (1978) Thalamocortical projections to the temporal and parietal association cortices in the rat. Neuroscience letters, 7, 121–125.

McNaughton, B.L., Mizumori, S.J., Barnes, C.A., Leonard, B.J., Marquis, M. & Green, E.J. (1994) Cortical representation of motion during unrestrained spatial navigation in the rat. Cereb Cortex, 4, 27–39.

Montero, V.M. (1993) Retinotopy of cortical connections between the striate cortex and extrastriate visual areas in the rat. Experimental brain research. Experimentelle Hirnforschung, 94, 1–15.

Morcos, A.S. & Harvey, C.D. (2016) History-dependent variability in population dynamics during evidence accumulation in cortex. Nature neuroscience, 19, 1672–1681.

Nitz, D.A. (2006) Tracking route progression in the posterior parietal cortex. Neuron, 49, 747–756.

Nitz, D.A. (2012) Spaces within spaces: rat parietal cortex neurons register position across three reference frames. Nature neuroscience, 15, 1365–1367.

Oh, S.W., Harris, J.A., Ng, L., Winslow, B., Cain, N., Mihalas, S., Wang, Q., Lau, C., Kuan, L., Henry, A.M., Mortrud, M.T., Ouellette, B., Nguyen, T.N., Sorensen, S.A., Slaughterbeck, C.R., Wakeman, W., Li, Y., Feng, D., Ho, A., Nicholas, E., Hirokawa, K.E., Bohn, P., Joines, K.M., Peng, H., Hawrylycz, M.J., Phillips, J.W., Hohmann, J.G., Wohnoutka, P., Gerfen, C.R., Koch, C., Bernard, A., Dang, C., Jones, A.R. & Zeng, H. (2014) A mesoscale connectome of the mouse brain. Nature, 508, 207–214.

Olcese, U., Iurilli, G. & Medini, P. (2013) Cellular and synaptic architecture of multisensory integration in the mouse neocortex. Neuron, 79, 579–593.

Olsen, G.M., Ohara, S., Iijima, T. & Witter, M.P. (2017) Parahippocampal and retrosplenial connections of rat posterior parietal cortex. Hippocampus, 27, 335–358.

Olsen, G.M. & Witter, M.P. (2016) Posterior parietal cortex of the rat: Architectural delineation and thalamic differentiation. The Journal of comparative neurology, 524, 3774–3809.

Olson, C.R. & Lawler, K. (1987) Cortical and subcortical afferent connections of a posterior division of feline area 7 (area 7p). The Journal of comparative neurology, 259, 13–30.

Padberg, J. & Krubitzer, L. (2006) Thalamocortical connections of anterior and posterior parietal cortical areas in New World titi monkeys. The Journal of comparative neurology, 497, 416–435.

Paxinos, G. & Franklin, K. (2012) Paxino’s and Franklin’s the Mouse Brain in Stereotaxic Coordinates. Academic Press.

Paxinos, G. & Watson, C. (2013) The Rat Brain in Stereotaxic Coordinates. Academic Press.

Raposo, D., Kaufman, M.T. & Churchland, A.K. (2014) A category-free neural population supports evolving demands during decision-making. Nature neuroscience, 17, 1784–1792.

Raposo, D., Sheppard, J.P., Schrater, P.R. & Churchland, A.K. (2012) Multisensory decision-making in rats and humans. J Neurosci, 32, 3726–3735.

Reep, R.L., Chandler, H.C., King, V. & Corwin, J.V. (1994) Rat posterior parietal cortex: topography of corticocortical and thalamic connections. Experimental brain research. Experimentelle Hirnforschung, 100, 67–84.

Remple, M.S., Reed, J.L., Stepniewska, I. & Kaas, J.H. (2006) Organization of frontoparietal cortex in the tree shrew (Tupaia belangeri). I. Architecture, microelectrode maps, and corticospinal connections. The Journal of comparative neurology, 497, 133–154.

Save, E. & Poucet, B. (2009) Role of the parietal cortex in long-term representation of spatial information in the rat. Neurobiol Learn Mem, 91, 172–178.

Schmahmann, J.D. & Pandya, D.N. (1990) Anatomical investigation of projections from thalamus to posterior parietal cortex in the rhesus monkey: a WGA-HRP and fluorescent tracer study. The Journal of comparative neurology, 295, 299–326.

Stepniewska, I., Cerkevich, C.M. & Kaas, J.H. (2016) Cortical Connections of the Caudal Portion of Posterior Parietal Cortex in Prosimian Galagos. Cereb Cortex, 26, 2753–2777.

Wang, Q. & Burkhalter, A. (2007) Area map of mouse visual cortex. The Journal of comparative neurology, 502, 339–357.

Wang, Q., Sporns, O. & Burkhalter, A. (2012) Network analysis of corticocortical connections reveals ventral and dorsal processing streams in mouse visual cortex. J Neurosci, 32, 4386–4399.

Whitlock, J.R., Pfuhl, G., Dagslott, N., Moser, M.B. & Moser, E.I. (2012) Functional split between parietal and entorhinal cortices in the rat. Neuron, 73, 789–802.

Whitlock, J.R., Sutherland, R.J., Witter, M.P., Moser, M.B. & Moser, E.I. (2008) Navigating from hippocampus to parietal cortex. Proc Natl Acad Sci U S A, 105, 14755–14762.

Whitlock, J.R. (2014) Navigating actions through the rodent parietal cortex. Front Hum Neurosci, 8, 293.

Wickersham, I.R., Lyon, D.C., Barnard, R.J., Mori, T., Finke, S., Conzelmann, K.K., Young, J.A. & Callaway, E.M. (2007) Monosynaptic restriction of transsynaptic tracing from single, genetically targeted neurons. Neuron, 53, 639–647.

Wilber, A.A., Clark, B.J., Demecha, A.J., Mesina, L., Vos, J.M. & McNaughton, B.L. (2014a) Cortical connectivity maps reveal anatomically distinct areas in the parietal cortex of the rat. Front Neural Circuits, 8, 146.

Wilber, A.A., Clark, B.J., Forster, T.C., Tatsuno, M. & McNaughton, B.L. (2014b) Interaction of egocentric and world-centered reference frames in the rat posterior parietal cortex. J Neurosci, 34, 5431–5446.

Wilber, A.A., Skelin, I., Wu, W. & McNaughton, B.L. (2017) Laminar Organization of Encoding and Memory Reactivation in the Parietal Cortex. Neuron, 95, 1406–1419 e1405.

Wise, S.P., Boussaoud, D., Johnson, P.B. & Caminiti, R. (1997) Premotor and parietal cortex: corticocortical connectivity and combinatorial computations. Annual review of neuroscience, 20, 25–42.

Zilles, K. (1985) The cortex of the rat: a stereotaxic atlas. Springer-Verlag Berlin Heidelberg.

Zingg, B., Hintiryan, H., Gou, L., Song, M.Y., Bay, M., Bienkowski, M.S., Foster, N.N., Yamashita, S., Bowman, I., Toga, A.W. & Dong, H.W. (2014) Neural networks of the mouse neocortex. Cell, 156, 1096–1111.

